# HIV-1 Vpr drives a tissue residency-like phenotype during selective infection of resting memory T cells

**DOI:** 10.1101/2021.01.25.428084

**Authors:** Ann-Kathrin Reuschl, Maitreyi Shivkumar, Dejan Mesner, Laura J. Pallett, José Afonso Guerra-Assunção, Rajhmun Madansein, Kaylesh J Dullabh, Alex Sigal, John P. Thornhill, Carolina Herrera, Sarah Fidler, Mahdad Noursadeghi, Mala K. Maini, Clare Jolly

## Abstract

Human immunodeficiency virus type 1 (HIV-1) replicates in CD4+ T cells leading to profound T cell loss, immunological dysfunction and AIDS. Determining how HIV-1 shapes the immunological niche in which it resides to create a permissive environment is central to informing efforts to limit pathogenesis, disturb viral reservoirs and achieve a cure. A key roadblock in understanding HIV-T cell interactions is the requirement to activate CD4+ T cells *in vitro* in order to make them permissive to infection. This dramatically alters T cell biology, obscuring native virus-host interactions. Here we show that HIV-1 cell-to-cell spread permits efficient and productive infection of resting CD4+ T cells without the need for prior activation. Infection is preferential for resting memory T cells, is observed with both CXCR4-tropic virus and CCR5-tropic transmitter-founder viruses and results in virus production and onward spreading infection. Strikingly, we find that HIV-1 infection of resting memory CD4+ T cells primes for induction of a tissue-resident memory (T_RM_)-like phenotype evidenced by upregulation of T_RM_ markers CD69/CXCR6 alongside co-expression of CD49a, PD-1, CD101 as well as transcription factor Blimp-1. Furthermore, we reveal that HIV-1 initiates a transcriptional program that overlaps with the core T_RM_ transcriptional signature. This reprograming depends on the HIV-1 accessory protein Vpr. We propose that HIV-1 infection drives a CD4+ T_RM_-phenotype potentially sequestering infected cells within tissues to support viral replication and persistence.

## Introduction

Resting primary CD4+ T cells cannot be efficiently infected by cell-free HIV-1 virions *in vitro* and require robust mitogenic stimulation to support viral replication^1–3^. This has led to the notion that T cell activation is necessary for HIV-1 replication and that resting T cells are not permissive for HIV-1 infection. However, mitogenic T cell activation *in vitro* results in wide-spread phenotypic and functional reprogramming which dominates changes to gene and protein expression^4–6^, concealing and altering authentic virus-host interactions. This presents a significant challenge for understanding the cellular response to HIV-1 infection and the consequences of the virus-host interaction for HIV-1 replication and persistence. While it is clear HIV-1 efficiently infects and replicates in activated T cells, the outcomes of the virus-host interaction with resting T cells has been reported to be cell death^7^ or latency^8^. However, previous data demonstrating that HIV-1 cell-to-cell spread is highly-efficient and drives widespread changes in protein phosphorylation status in both infected and target cells^9–13^ suggested that cell-to-cell spread may overcome the barrier to productive infection of resting T cells. Here, we comprehensively show for the first time that HIV-1 exploits cell-to-cell spread to efficiently infect resting memory CD4+ T cells and have used this to uncover a hithertho unknown consequence of HIV-1 infection for T cell reprogramming driven by the accessory protein Vpr.

## Results

To test whether cell-to-cell spread allows for productive infection of resting T cells, HIV-1 infected Jurkat T cells (Fig. 1a) or primary CD4+ T cells (Fig. 1b) were co-cultured with uninfected resting CD4+ T cells (Extended data Fig. 1a-e). We confirmed that CD4+ T cells isolated from peripheral blood display a resting phenotype by staining for Ki67, CD69, HLA-DR and MCM2. (Extended data Fig. 1c-e). Infection of resting target cells in the absence of mitogenic or cytokine activation was measured. Direct co-culture of infected and uninfected cells resulted in significant levels of HIV-1 infection of resting CD4+ target T cells (Gag+) measured by intracellular flow cytometry staining (Fig. 1a, b and Extended data Fig. 1f). By contrast, resting CD4+T cells were not infected (<1%) when cell-cell contact was prevented by separating the two cell populations by a transwell (Fig. 1a,b), a condition that only allows for cell-free infection. As expected, mitogenic activation of target T cells made them more permissive to cell-free HIV-1 but, as previously shown, infection was still substantially boosted by direct co-culture allowing for additional cell-to-cell spread (Extended data Fig.1f-h) ^9, 11, 12^. Infection of resting CD4+ target T cells was preferentially detected in CD45RA-ve resting memory T cell populations rather than CD45RA+ naïve T cells, which are both abundant in peripheral blood (Fig. 1c-e, Extended Fig. 1i). Co-staining for CD62L confirmed that the infected CD45RA+ve cells were mainly naïve rather than T_EMRA_ (Extended data Fig. 1j,k)^14^. The preferential infection of CD45RA-ve memory T cell populations rather than the CD45RA+ve naïve population, is in agreement with HIV-1 being predominantly detected in memory CD4+ T cells *in vivo* ^15–17^. This was not due to competition between naïve and memory cells because the same effect was observed when CD45RA+ve and CD45RA-ve resting CD4+ target T cells were separated prior to cell-to-cell infection (Fig. 1f and Extended data Fig. 2a). Cell-to-cell infection of resting memory T cells was observed with CXCR4-tropic strain NL4.3, and CCR5-tropic viruses NL4.3 BaL and two transmitter-founder (T/F) primary isolates (CH040 and CH077) (Fig. 1g-i) demonstrating that selective permissivity was not unique to a particular virus or receptor tropism. Preventing viral entry with fusion inhibitor (T20), or blocking reverse transcription (Efavirenz), inhibited the appearance of Gag+ve or GFP+ve cells (Fig. 1j-l and Extended data Fig. 2b,c) demonstrating that this signal reflects productive infection and not simply virus capture^9, 11–13^. Consistent with infection, we also observed downregulation of CD4 expression on target cells (Fig. 1m and Extended data Fig. 2d) that was most pronounced in the resting memory T cell population. Importantly, flow-sorting and culturing HIV-1 infected resting target T cells showed that these cells support viral replication by producing new virus and transmitting infection to new target cells (Fig. 1n-p and Extended data Fig. 2e), further demonstrating productive infection.

**Fig. 1.**
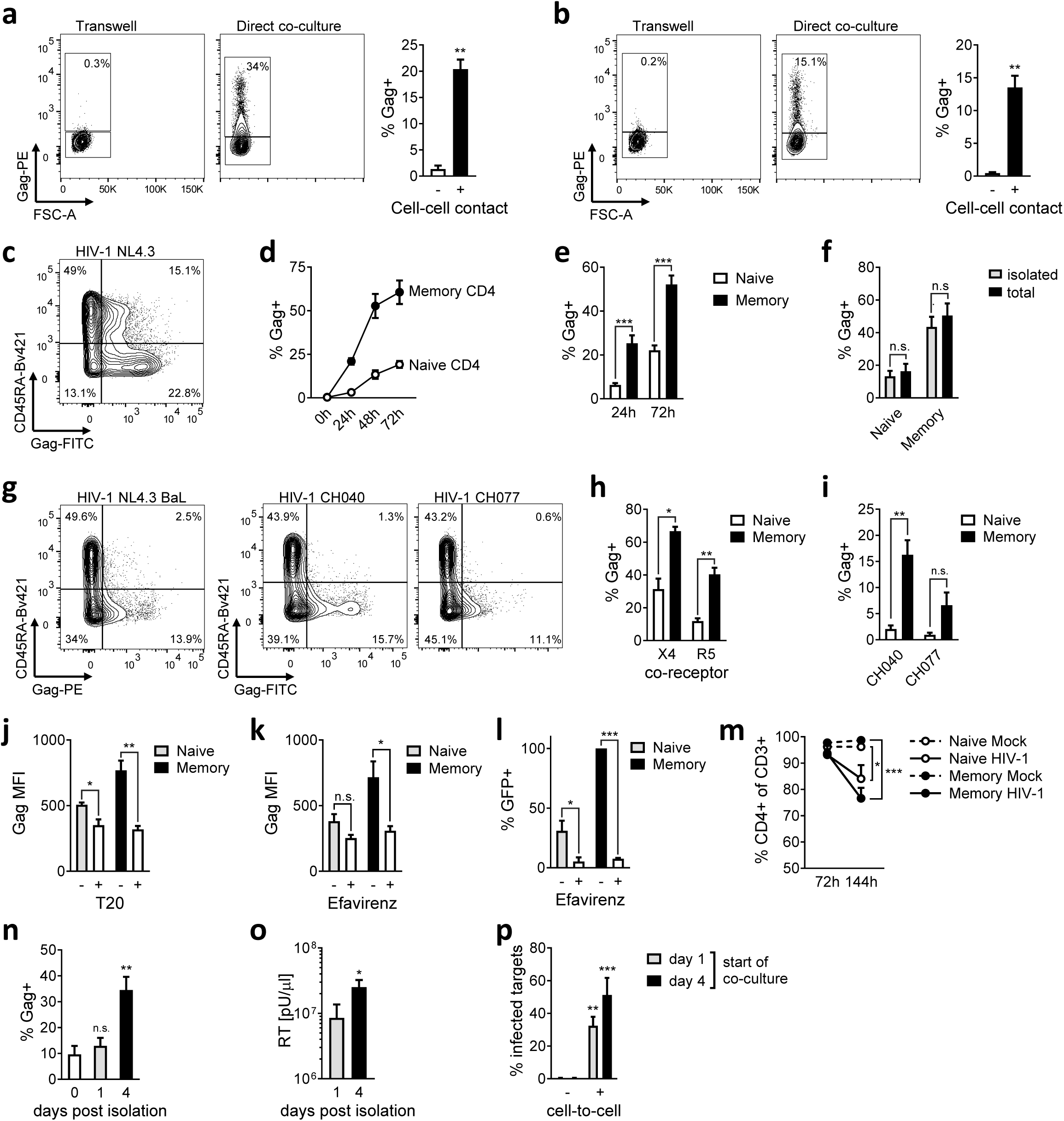
HIV-1 exploits cell-to-cell spread to preferentially infect resting memory CD4+ T cells. HIV-1 NL4.3 infected (**a**) Jurkat or (**b**) mitogenically-activated primary CD4+ donor T cells co-cultured with resting primary CD4+ target T cells separated by a 0.4μm transwell (cell-free) or direct co-culture (cell-cell). Target cell infection was measured by intracellular staining for HIV-1 Gag protein. Representative flow cytometry plots are shown. Bar graphs show mean of independent experiments (n=4). (**c,d**) Cell-to-cell spread from activated primary donor CD4+ T cells to resting primary target CD4+ T cells preferentially infects CD45RA-memory CD4+ T cells. A representative flow cytometry plot and quantification is shown (n=4). (**e**) Quantification of infection performed as in (**c**) (n=11). (**f**) HIV-1 infection of target CD4+ T cells as part of the total resting CD4+ T cell population (total) compared to pre-isolated naive and memory CD4+ target T cells (isolated) (n=9). (**g**) Representative flow cytometry plots of cell-to-cell infection of resting CD4+ T cells with CCR5-tropic HIV-1 NL4.3 BaL and transmitter founder viruses HIV-1 CH040 and CH077 as performed in (**c**). (**h**) Quantification of infection of CXCR4 (X4) and CCR5 (R5)-tropic viruses (n=4) and (**i**) transmitter/founder viruses HIV-1 CH040 and CH077 (n=7). (**j**) Cell-to-cell infection of resting CD4+ T cells is reduced by the HIV-1 fusion-inhibitor T20 (n=6) and (**k**) the reverse transciptase inhibitor Efavirenz (n=6) measured by intracellular Gag staining (MFI) or (**l**) HIV-1 LTR- driven GFP-reporter gene expression (n=4). (**m**) HIV-1 infection downregulates CD4 expression. Shown are the percentage of CD4+ cells of the total CD3+ target cell population (n=6). Resting CD4+ memory T cells were isolated after 72h of cell-to-cell spread by FACS sorting and cultured for 4 days. HIV-1 infection was measured by intracellular Gag staining (**n**) and virus release measured by culture supernatant RT activity (**o**) (n=5-7). T cells from (**n**) were recovered at day 1 or 4 post isolation and cultured with uninfected eFluor450+ target Jurkat T cells. Infection of Jurkat T cells was measured after 72h (**p**) (n=3). Data are the mean±SEM. Paired two-tailed *t*-test or one-way ANOVA with Bonferroni post-test were used. For (**m**), median+IQR is shown and Friedman test with Dunn’s post-test was used. For (**o**), unpaired one-tailed *t*-test was used. *, p<0.05; **, p<0.01; ***, p<0.001; n.s., not significant.

We confirmed that HIV-1+ target T cells infected by cell-to-cell spread maintained their resting phenotype and did not upregulate Ki67 or MCM2, two markers of cell-cycle progression (Extended data Fig. 3a,b), showing these cells were not simply being activated and driven into cell cycle, either by infection or bystander effects during co-culture. Intriguingly, expression of CD69 on HIV-1 infected resting memory target T cells was significantly increased compared to mock-treated target T cells (Fig. 2a,b). This was not due to preferential infection of a minor pre-existing population of CD69+ve CD4+ T cells in blood (Extended data Fig. 1c) because flow cytometry sorting of CD69-ve CD4+ T cells (prior to co-culture with HIV-1 infected donor T cells) showed *de novo* upregulation of CD69 on the newly-infected resting memory target cells (Extended data Fig.3c,d). While CD69 is classically thought of as a marker of early T cell activation, expression can occur independently of cell cycle progression and T cell activation^18–20^. Consistent with this we did not detect T cell activation concomitant with CD69 upregulation in HIV-1 infected resting CD4+ memory T cells since CD69+ve cells remained HLA-DR-negative (Extended data Fig.3e).

**Fig. 2.**
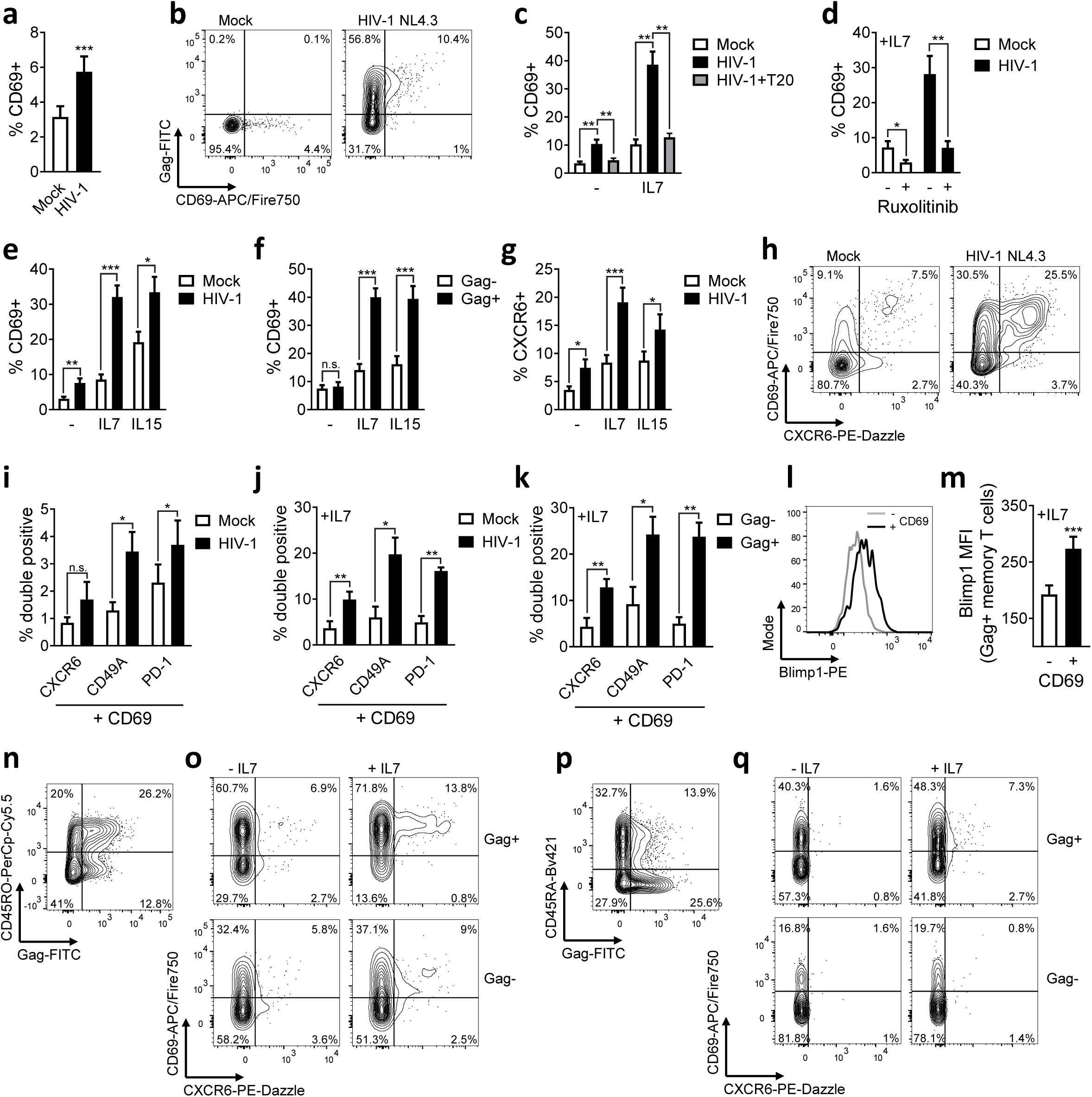
HIV-1 infection induces a T_RM_-like phenotype in resting memory CD4+ T cells. (**a**) CD69 expression on resting memory CD4+ target T cells following co-culture with HIV-1 infected primary donor T cells or uninfected donor T cells (mock) (n=17). (**b**) Representative flow cytometry plots from (**a**). (**c**) CD69 expression on infected resting memory CD4+ T cells ± IL7 and T20 (n=7). (**d**) CD69 expression on infected resting memory CD4+ T cells ± IL7 and Ruxilitinib (n=8). (**e**) CD69 expression on infected resting memory CD4+ T cells in response to IL7 and IL15 (n=11). (**f**) CD69 expression on infected Gag+ resting memory CD4+ T cells and uninfected Gag-bystander cells in response to IL7 and IL15 (n=11). (**g**) CXCR6 surface expression from (**f**) (n=11). (**h**) Representative flow cytometry plots of CD69 and CXCR6 co-expression. (**i**) Co-expression of CD69 with CXCR6, CD49A or PD-1 on infected resting memory CD4+ T cells (n=5-7). (**j**) As for (**i**) in the presence of IL7 (n=4-7). (**k**) As for (**i**) comparing infected Gag+ memory CD4+ T cells and uninfected Gag-bystander cells. (**l**) and (**m**) Blimp-1 expression in CD69+ infected resting memory CD4+ T cells and infected CD69- cells in response to IL7 (n=8). (**n**) Total lymphocytes from cellularised tonsils co-cultured with HIV-1 infected Jurkat T cells. Infection of resting CD4+ T cells (live CD3+/CD8-/Ki67- lymphocytes) shown as CD45RO vs Gag. (**o**) CD69 and CXCR6 co-expression on infected Gag+ and uninfected Gag-tonsil resting memory CD4+ T cells ± IL7. (**p**) Total lymphocytes from mediastinal lymph nodes co-cultured with HIV-1 infected activated autologous LN-derived lymphocytes. Infection of resting CD4+ T cells (live CD3+/CD8-/Ki67- lymphocytes) shown as CD45RA vs Gag. (**q**) CD69 and CXCR6 co-expression on infected Gag+ and uninfected Gag-lymph node resting memory CD4+ T cells ±IL7. Data are the mean±SEM. Paired two-tailed *t*-test or one-way ANOVA with Bonferroni or Dunnett’s post-test were used. *p<0.05; **, p<0.01; ***, p<0.001; n.s., not significant.

Functionally, CD69 is crucial for T cell retention in tissues by interfering with S1P receptor mediated egress and has been identified as a hallmark of tissue-resident memory (T_RM_) T cells^21^. Recently it has been demonstrated that although T_RM_ cells are largely absent in peripheral blood (Extended data Fig. 3f), precursor cells poised to adopt a T_RM_ phenotype are present in circulation^22, 23^. Interestingly, while HIV-1 infection alone was associated with upregulation of CD69, exposure of HIV-1 infected resting memory T cells to the homeostatic T cell cytokine IL7 further boosted CD69 upregulation 4-fold, as compared to HIV-1 alone or IL7 alone (Fig. 2c and Extended data Fig. 3j). IL7 secreted by stromal cells is required for long-term maintenance of CD4+ T_RM_ cells^24–26^. Although IL7 can enhance HIV-1 infection of T cells^27^ (Extended data Fig. 3g), infection of resting memory T cells mediated by cell-to-cell spread does not require IL7 (Fig.1) and IL7 increased CD69 expression on infected cells even when added 48h post-infection (Extended data Fig. 3j). HIV-1 induced CD69 upregulation was completely abrogated by suppressing infection with the fusion inhibitor T20 (Fig. 2c) or by treating cells with Ruxolitinib that blocks IL7-mediated JAK signalling (Fig. 2d) demonstrating that the enhanced CD69 induction requires both infection and cytokine signalling. Similar infection-driven enhancement of CD69 expression was observed in response to γ_c_-chain cytokine IL15 (Fig. 2e,f) but not IL12 or TGF-β (Extended data Fig.3h,i). Consistent with the hypothesis that HIV-1 infection induces a CD4+ T_RM_-like phenotypic signature, resting memory T cells also upregulated the T_RM_ marker CXCR6^21^ during HIV-1 infection, (Fig. 2g,h and Extended data Fig. 3k,n), as well as increasing the population of CD69+ve cells co-expressing CD49a, PD-1 and CD101 that are also associated with the core T_RM_ signature^21^ (Fig. 2i, j and Extended data Fig. 3l). As expected for T_RM_s, we saw no upregulation of CX3CR1 expression (Extended data Fig. 3o) and no transcriptional upregulation of *S1PR1* or *KLF2* mRNA (Extended data Fig. 5i). Critically, induction of T_RM_-associated markers did not occur in uninfected Gag-ve bystander cells (Fig. 2k and Extended data Fig.3m,n). By contrast to CD8+ T_RM_ cells, CD103 was barely detectable and not upregulated (Extended data Fig.3p,q), consistent with the observation of limited CD103 expression on CD4+ T cells^21^. Induction of CD69 expression was also concomitant with upregulation of the T_RM_-associated transcription factor Blimp-1^28–30^ (Fig. 2l,m). Similar upregulation of T_RM_-associated markers were also observed when unstimulated CD4+ T cells from tonsil (Fig. 2n,o) or mediastinal lymph nodes (Fig. 2p,q) were infected with HIV-1 via cell-to-cell spread and exposed to IL7 (Fig. 2o,q), demonstrating that induction of TRM-like phenotype also occurs in tissue-derived T cells following HIV-1 infection. Taken together, these data suggest that HIV-1 infection of resting memory CD4+ T cells reprogrammes cells by upregulating expression of T_RM_-associated marker proteins, and thus induces a phenotype characteristic of tissue residency.

HIV-1 expresses four accessory proteins, Vif, Vpu, Vpr and Nef which directly and indirectly manipulate host cell factors to facilitate efficient viral replication *in vivo* and drive pathogenesis^31^. Co-culture of resting target T cells with donor T cells infected with HIV-1 accessory protein mutants showed that deletion of Vpr (HIV-1 ΔVpr) resulted in a complete abrogation of the T_RM_-like phenotype as evidenced by a lack of CD69 upregulation, and no detection of the CD69+/CXCR6+/CD49a+ triple-positive T_RM_-like memory population (Fig. 3a,b,d, Extended data Fig.4 and 5a-e). By contrast, deletion of Vpu or Nef did not affect HIV-1 induction of T_RM_-markers on infected resting T cells (Fig. 3a,b, Extended data Fig. 4 and Fig. 5a-e). HIV-1 ΔVif could not be tested because Vif is required to antagonise APOBEC3-mediated viral restriction and allow infection^32^. HIV-1 Vpr is not required for infection of T cells *in vitro*^33, 34^ and concordantly, loss of T_RM_-marker protein induction was not due to lack of infection of target cells by HIV-1 ΔVpr virus, nor reduced Gag expression (Fig. 3c and Extended data Fig. 5f-h). Critically, Vpr was also required for induction of CD69 expression observed at the mRNA level, as well as induction of *CXCR6* and Blimp1 (*PRDM1*) mRNA (Fig. 3e). As expected, there was no upregulation of *S1PR1* or *KLF2* mRNA by either HIV-1 WT or ΔVpr virus (Extended data Fig.5i), consistent with their suppression under conditions of T_RM_ induction^21^. HIV-1 dependent upregulation of CD69 surface expression was inhibited by Ruxolitinib treatment (Fig. 2d) in a Vpr-dependent manner (Extended data Fig. 5j). Furthermore, HIV-1 infection downregulated the IL7 receptor alpha-subunit (CD127) from the cell surface (Extended data Fig 5k) and transcriptionally (fold change=0.693, adjusted p-value=5.89E-08, Supplementary table 1), indicative of activation of this pathway^35^ by HIV-1. This was accompanied by a significant increase in the intracellular levels of phosphorylated transcription factor STAT5 (Extended data Fig. 5j and k). These data are consistent with Vpr manipulating cellular signalling pathways to drive induction of a T_RM_-like phenotype in response to IL7. Vpr-dependent T_RM_-like induction was also accompanied by spontaneous production of IFNγ by infected CD4+ memory T cells (Fig. 3f), in line with the increased capacity of T_RM_ to produce this cytokine^21, 36^. Notably, Vpr-mediated induction of a T_RM_-like phenotype was not inhibited by the integrase inhibitor Raltegravir (Fig. 3g, h) that potently suppressed HIV-1 integration into resting memory CD4+ T cells infected by cell-to-cell spread (Fig. 3i), demonstrating that the T_RM_- like phenotype does not require integration. Of note, the presence of integrated provirus in resting CD4+ T cells (less than or equal to 1 provirus per cell Fig. 3i) further supports the observation that cell-to-cell spread results in resting CD4+ T cell infection (Fig. 1). Collectively, these data identify Vpr as the viral determinant required for upregulation of T_RM_-associated proteins on HIV-1 infected resting CD4+ T cells.

**Fig. 3.**
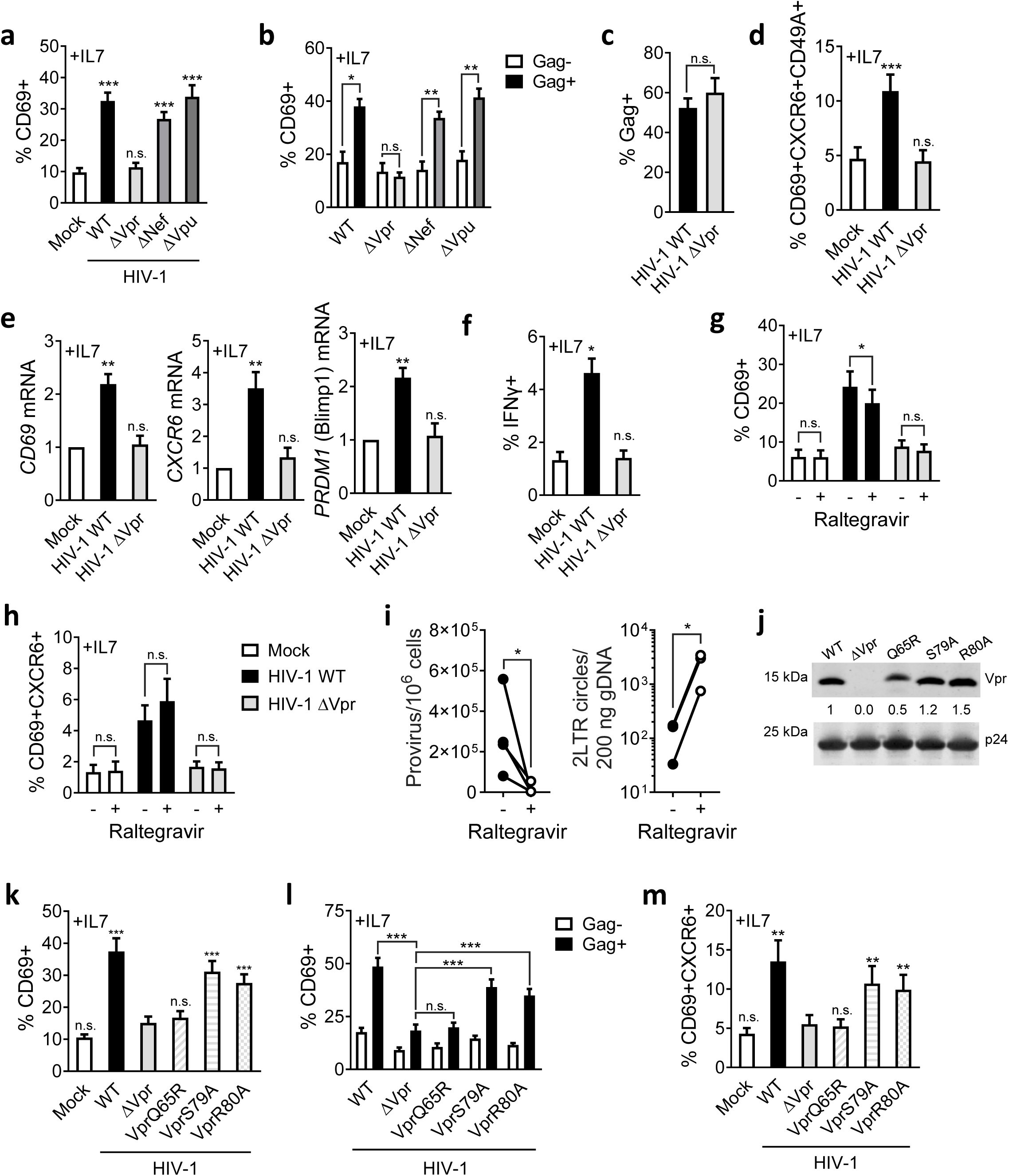
Vpr drives HIV-1-induced T_RM_-induction in resting memory CD4+ T cells. Resting memory CD4+ T cells were co-cultured with HIV-1 infected primary CD4+ T cells infected with HIV-1 WT or mutant viruses, or uninfected donor cells (mock). (**a**) CD69 upregulation in response to IL7 compared to mock (n=9). (**b**) CD69 expression on HIV-1 infected Gag+ resting memory CD4+ T cells compared to uninfected Gag-bystander cells (n=9). (**c**) Quantification of cell-to-cell spread of HIV-1 WT and ΔVpr to resting memory CD4+ T cells (n=9). (**d**) CD69/CXCR6/CD49a co-expression on resting memory CD4+ T cells infected with HIV-1 WT or ΔVpr (n=9). (**e**) *CD69*, *CXCR6* and *PRDM1* (Blimp1) mRNA levels from FACS sorted infected resting memory CD4+ T cells. Fold change relative to unifected (mock) is shown (n=5). (**f**) IFNg expression by HIV-1 infected resting memory CD4+ T cells at 72h in response to IL7 (n=3). (**g**) CD69 and (**h**) CD69/CXCR6 co-expression in response to IL7 in the presence of integrase inhibitor Raltegravir (n=6). (**i**) Quantification of integrated provirus and 2LTR circles in FACS sorted target CD4+ memory T cells after 72h of cell-to-cell spread in the presence or absence of Raltegravir. (**j**) Western blot showing Vpr packaging into HIV-1 WT and Vpr-mutant virions. (**k**) CD69 upregulation in response to IL7 on resting memory CD4+ T cells infected with HIV-1 WT, ΔVpr or Vpr mutants (n=9). (**l**) as for (k) showing CD69 expression on HIV-1 infected Gag+ memory T cells compared to uninfected Gag-bystander cells. (**m**) Co-expression of CD69 and CXCR6 from (k) (n=9). Data are the mean±SEM. Paired two-tailed *t*-test or one-way ANOVA with Bonferroni or Dunnett’s post-test were used. 2LTR circles (i) were compared by unpaired one-tailed *t*-test. *, p<0.05; **, p<0.01; ***, p<0.001; n.s., not significant.

Vpr is a multifunctional protein that is packaged into viral particles and is present during the early stages of infection where it plays an important, but as yet poorly-defined role in HIV-1 pathogenesis. Amongst the best-defined functions of Vpr are its ability to a) bind the Cul4A-DDB1 (DCAF1) complex (leading to an interaction with the ubiquitinylation and proteasomal machinery); b) induce G2/M cell-cycle arrest and; c) drive apoptosis in infected cells^37–41^. We abrogated these functions individually by introducing the mutations Q65R, S79A or R80A into Vpr in the context of virus and confirmed that each Vpr mutant is packaged into virions (Fig. 3j). Co-culture of resting memory target T cells with HIV-1+ T cells infected with different Vpr mutants revealed that the Vpr mutants S79A or R80A (inhibits cell cycle arrest) behaved similarly to WT virus and did not induce CD69 and CXCR6 upregulation (Fig. 3k-m). By contrast, Q65R (that is most closely associated with loss of DCAF1 binding) was unable to induce a T_RM_-like phenotype following infection of target cells, behaving like ΔVpr virus in these experiments (Fig. 3k-m).

Next we performed transcriptional profiling by RNA-Seq analysis of flow-sorted cells infected with HIV-1 WT or ΔVpr virus by cell-to-cell spread. Figure 4 shows that HIV-1 infection induced widespread changes to gene expression in resting memory T cells when compared to uninfected cells (232 differentially expressed genes (DEG), fold change >1.2, adjusted p-value<0.01) (Fig. 4). Hierarchial clustering and principal component analysis revealed that the gene expression patterns in response to HIV-1 WT infection were clearly distinct from those induced following infection with ΔVpr virus (Fig. 4 a-c, e,f) demonstrating that *Vpr*-deletion suppresses the global transcriptional response to HIV-1 infection. Specifically, infection with ΔVpr virus resulted in only 13 genes showing statistically significantly changes compared to uninfected cells, by contrast to 232 genes for HIV-1 WT virus (Supplementary table 1 and 3, Fig. 4e). In fact, much of the transcriptional response to HIV-1 infection was regulated by Vpr, as evidenced by changes in DEG in the presence and absence of Vpr (Fig. 4f). The requirement for Vpr in driving many of the changes to DEG was also observed in response to IL7 (Fig. 4d, Supplementary table 2 and 4, and Extended data Fig. 6a). Consistent with Vpr manipulating the T cell response to HIV-1 infection (Supplementary table 5 and 6), gene-set enrichment analysis (GSEA) (Fig. 4g) and Ingenuity Pathway Analysis (IPA) (Fig. 4h and i) revealed enrichment of numerous cellular signalling pathways following HIV-1 infection that appeared Vpr-dependent, most notably pathways associated with cytokine and inflammatory responses as well as immune signalling (Fig. 4g, Extended data Fig. 6 and Supplementary table 7 and 8). This was further evidenced by Upstream Regulator analysis that showed significant enrichment for genes associated with cytokine signalling and transcriptional regulators that were again largely Vpr-dependent (Fig. 4j and k, and Extended data Fig. 6). Taken together, these data reveal that HIV-1 induces dramatic reprogramming during infection of resting memory CD4+ T cells that is driven largely by Vpr.

**Fig. 4.**
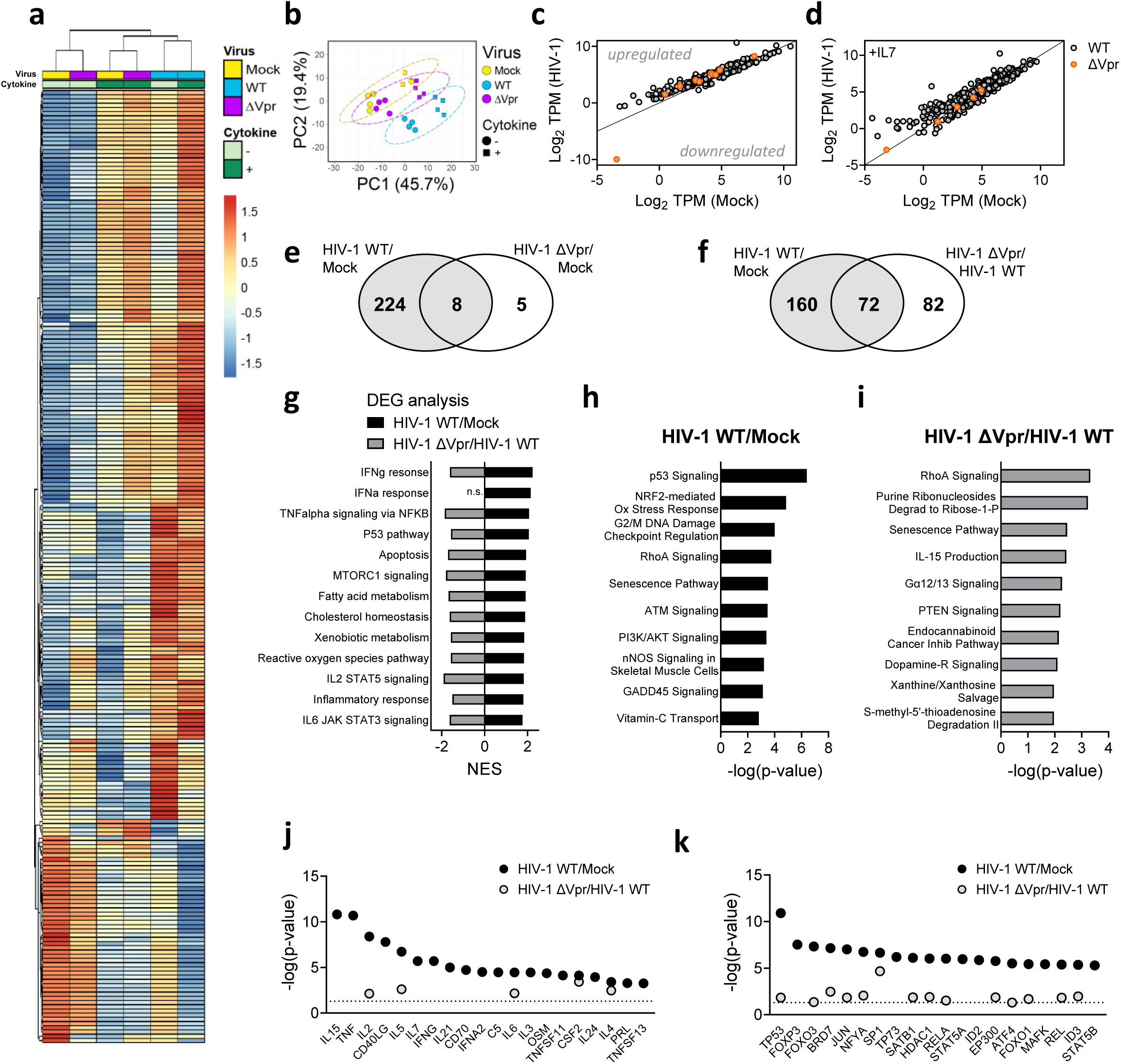
Transcriptional profiling of HIV-1 infected resting memory CD4+ T cells. (**a**) Heatmap showing hierarchical clustering of 226 differentially expressed genes (DEG) of infected (HIV-1 WT) over uninfected (Mock) resting memory CD4+ T cells (adjusted p-value < 0.01, Fold change +/- 1.2). Mean log2 TPM of 4 biological repeats are shown. Cytokine indicates presence or absence of IL7. Virus indicates infection with HIV-1 WT, HIV-1 ΔVpr or uninfected (Mock) conditions. (**b**) Principal component analysis (PCA) of (a), with ellipses indicating 95% CI. (**c**) and (**d**) show scatter plots of mean log2 TPMs of DEGs from HIV-1 WT/Mock (grey circles) or HIV-1 ΔVpr/Mock (orange circles) in the absence (c) or presence (d) of IL7 (adjusted p-value < 0.01, Fold change +/- 1.2). Lines indicate line of identity (LOD). Genes above or below LOD are up or downregulated, respectively. (**e**) and (**f**) Venn diagrams showing overlap of DEGs comparing expression profiles of HIV-1 WT/Mock with (e) HIV-1 ΔVpr/\Mock or (f) HIV-1 ΔVpr/HIV-1 WT. (**g**) GSEA was performed on expression profiles comparing HIV-1 WT / Mock (black) or HIV-1 ΔVpr/HIV-1 WT (grey). Normalised enrichment scores are shown for significantly enriched Hallmark gene sets are shown (FDR q-value<0.05 and NES>1.75). (**h**) and (**i**) top ten significantly enriched canonical pathways predicted by ingenuity pathway (IPA) analysis of DEGs (h) HIV-1 WT/Mock or (i) HIV-1 ΔVpr/HIV-1 WT (adjusted p-value<0.05). (**j**) Cytokines and (**k**) transcription regulators predicted to be upstream regulators by IPA of gene expression signatures HIV-1 WT/Mock (black) or HIV-1 ΔVpr/Mock (grey), line indicates p=0.05.

Tissue residency of T cells has been associated with a 31-gene core transcriptional signature^21^. We took advantage of this dataset and our RNAseq analysis to determine whether this core T_RM_ signature was enriched in our transcriptome of HIV-1 infected resting memory T cells. Hierarchical clustering of our data, compared to the core signature of T_RM_ cells (CD69+ve T cells isolated from human lung and spleen)^21^ from Kumar et al, showed that HIV-1 infected memory T cells exposed to IL7 grouped distinctly and clustered with CD69+ve T_RM_ cells (Fig. 5a, Supplementary table 9) distinct from non-T_RM_ T cells (CD69-ve T cells isolated from tissue and blood from Kumar et al). We further corroborated this finding by calculating a T_RM_ enrichment score based on the published core transcriptional signature, which showed that HIV-1 infected resting memory T cells (both +/-IL7) harbour a higher T_RM_ signature score, significanly different from non-T_RM_ T cells, approximating more closely to that of *bona fide* T_RM_ cells (Fig. 5b). Critically, this was Vpr-dependent, with mock and ΔVpr infected cells showing an enrichment score that was not statistically different to non-T_RM_. Taken together, these data demonstrate that HIV-1 induces both a phenotypic and transcriptional T_RM_-like signature in resting T cells via the accessory protein Vpr.

**Fig. 5.**
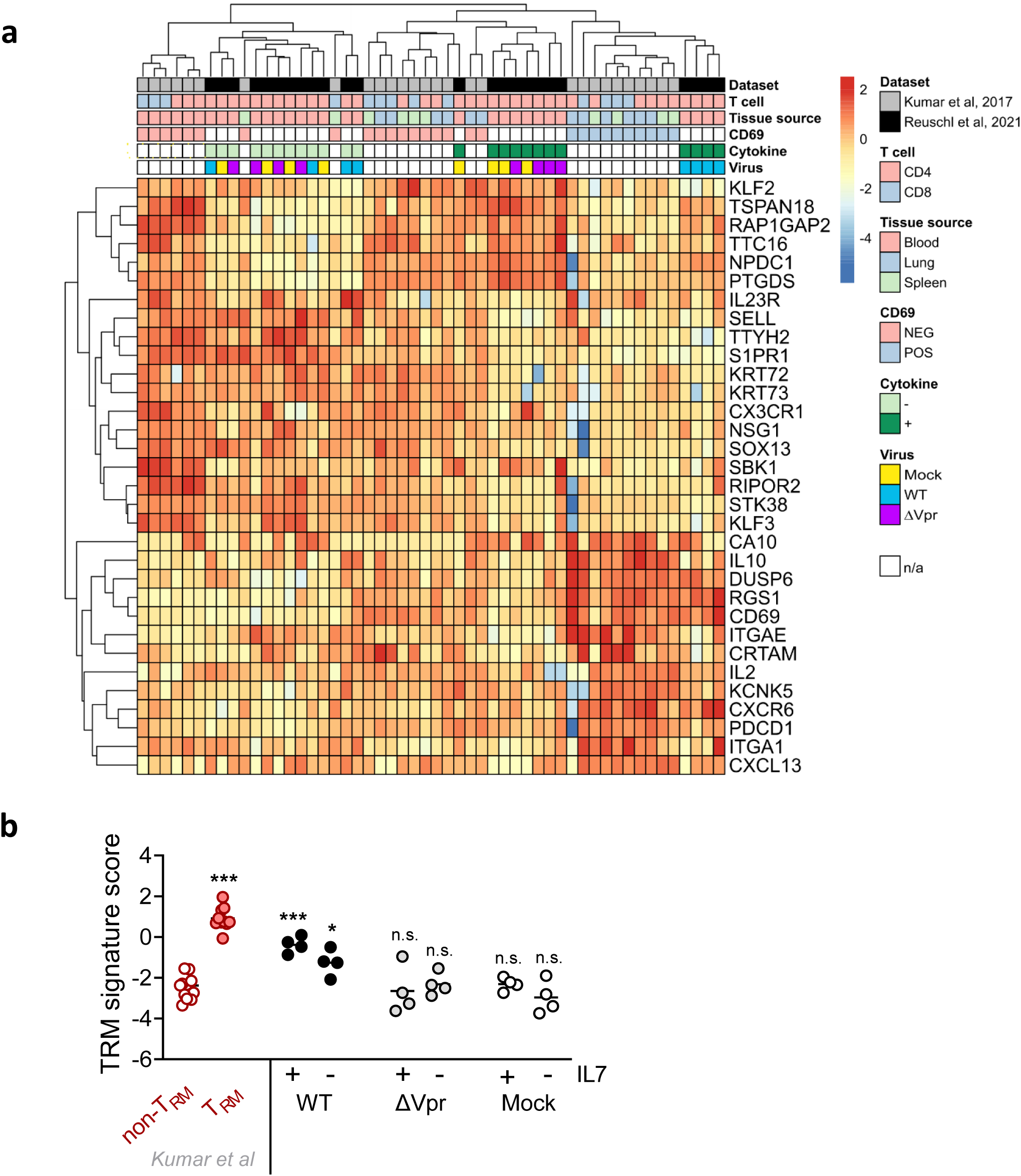
Vpr drives a T_RM_-like transcriptomic programme in HIV-1 infected resting memory CD4+ T cells. (**a**) Heatmap showing hierarchical clustering based on a T_RM_ core gene expression signature^21^ that was performed to compare transcriptional profiles of HIV-1 infected resting memory CD4 T cells (Reuschl et al, 2021) (Mock, HIV-1 WT, HIV-1 ΔVpr) with previously described gene expression profiles (Kumar et al, 2017) of T_RM_ (CD69 POS), non-T_RM_ (CD69 NEG) tissue-derived T cells (lung and spleen) and blood-derived CD69- (CD69 NEG) T cells. Cytokine indicates presence or absence of IL7. Virus indicates infection with HIV-1 WT, HIV-1 ΔVpr or uninfected (Mock) conditions. n/a, not applicable. (**b**) shows the T_RM_ signature score for the indicated conditions calculated based on (a). Subsets from^21^ are indicated in red, shown are CD4+ or CD8+ T cells from lungs or spleens. T_RM_+, CD69+ T cells; T_RM_-, CD69- T cells. T_RM_ signature scores for resting CD4 memory T cells infected or uninfected are shown in the presence or absence of IL7. Means are shown.

## Discussion

Our discovery that resting CD4+ T cells can be productively infected by cell-to-cell spread, allowing for viral integration, replication and dissemination, transforms our ability to determine how T cells respond to and suppport HIV-1 replication without confounding activation-induced changes. Recently, it has been reported that cell-to-cell spread can also facilitate latent infection of resting T cells without productive infection^8^. Together, these two studies highlight the distinct advantages of co-culture models (that do not require mitogenic, experimental stimulation of T cells to drive infection) to study native HIV-1-host cell interactions and the cellular response to infection.

Here we have employed our co-culture model to show that HIV-1 infection of resting memory T cells induces a T_RM_-like phenotype evidenced by upregulation and co-expression of T_RM_-associated markers on infected cells and induction of a core T_RM_ transcriptional signature. HIV-1 establishes cellular and tissue reservoirs (both active and latent) that ultimately prevent cure with antiretroviral therapy. Importantly, T_RM_ cells are long-lived and are thought to be largely confined to tissue^42^ providing an alternate model for a tissue-associated reservoir driven by the virus itself. Our results suggest that HIV-1 persistence and the establishment of tissue reservoirs may be driven, in part, through direct viral induction of a T_RM_-like phenotype via transcriptional reprogramming. Recently, TRM in cervical tissue were found to be preferentially infected by HIV-1 and can harbour an HIV-1 reservoir *in vivo*^43^. The relative contribution of pre-existing versus HIV-1 induced T_RM_ cells to viral reservoirs and their relative abundance in different anatomical compartments *in vivo* remains to be quantified, but we expect T_RM_ cells harbouring virus to be important contributors to viral persistence. In light of these findings it is possible that HIV-1 infected cells circulating in peripheral blood may in fact represent cells that have failed to become part of the tissue reservoir, leading to an underestimation of the true viral burden. Having shown that HIV-1 infection of resting T cells by cell-to-cell spread results in productive infection, we hypothesise that induction of a T_RM_-like phenotype in infected cells may also play additional roles in establishing and maintaining viral reservoirs by sequestering infected cells in tissue sites where susceptible target T cells are in abundance, thus supporting localised viral replication. Indeed we have shown that infected resting memory T cells support spreading infection to disseminate virus. Given the importance of T_RM_ cells as a population that are increasingly recognised to be critical in providing localised immunity and immunosurveillance^20, 30^, future work should focus on understanding the contribution of HIV-1 induced T_RM_-like cells in pathogenesis and persistence.

It is now emerging that committed T_RM_ precursors, imprinted with the capacity to become mature T_RM_ pre-exist in blood and that when exposed to the appropriate cues in tissues or *ex vivo* can become tissue-resident^22, 23^. Thus the ontogeny, derivation and maintenance of T_RM_ cells appears more complex that initially appreciated. Our discovery that HIV-1 induces a T_RM_-like phenotype in CD4+ T cells provides an opportunity to gain new understanding of mechanisms behind CD4+ T_RM_ induction and maintainance.

We found that HIV-1 infection of resting memory T cells was associated with striking transcriptional reprograming that was licenced by Vpr, thus identifting a novel function for this enigmatic HIV accessory protein. Notably, HIV-1 induction of a T_RM_-like phenotype via Vpr was accompanied by induction of a T_RM_ transcriptional signature that aligned closely with a published core T_RM_ signature^21^. Vpr-deletion not only abolished induction of this T_RM_ signature, but also many HIV-1 induced changes to gene expression following infection of resting T cells. This is in keeping with wide-spread proteome remodelling by Vpr in activated T cells^44^, but suggests that these changes may be driven in part by a hitherto unappreciated role for Vpr in modulating the host cell gene expression profile. Whether this reflects wide-spread epigenetic changes mediated by Vpr or manipulation of key upstream regulators remains to be determined. Our data show HIV-1 Vpr modulates T cell responsiveness to external stimuli by manipulation of immune signalling pathways, including innate and inflammatory responses. This is particularly intriguing and suggests HIV-1 manipulates immune signalling pathways to benefit the virus, in this case by priming resting memory T cells for T_RM_-like induction. Vpr-mediated induction of T_RM_-like phenotype was dependent on residue Q65 and Vpr is reported to drive wide-spread remodelling of the cellular proteome via its recruitment of DCAF1 through Q65^44^. Whether this requirement for Q65 in induction of a T_RM_-like phenotype is DCAF1-dependent or not remains unclear because DCAF1 knockdown in primary T cells made cells hyper-responsive to HIV-1 induced cell death (Extended data Fig. 7). Thus we cannot at present formally exclude other functions of Vpr Q65 in the process of T_RM_-induction.

Notably, a rare case of laboratory-derived infection with Vpr-defective HIV-1 was characterised by markedly delayed seroconversion, suppressed viremia and normal CD4+ T cell counts^45^, consistent with reduced pathogenesis and failure to establish and maintain a significantly large tissue reservoir. We envisage therapeutic targeting of Vpr to manipulate persistence and pathogenesis. In order to achieve an HIV-1 cure it is essential to understand the nature and establishment of HIV-1 reservoirs and how to manipulate them. By demonstrating that HIV-1 infection drives a T_RM_-like phenotype during infection of resting memory T cells we have taken a significant step towards this to help accelerate the quest for an HIV-1 cure.

## Methods

### Cells

Peripheral blood mononuclear cells (PBMC) were isolated from buffy coats from healthy donors (UK NHS Blood and Transplant Service) by density centrifugation using FicollPaque Plus (GE Life Sciences) and cryopreserved in 10% DMSO (Sigma-Aldrich) in 90% FBS (LabTech). Resting CD4+ T cells were isolated from total PBMCs by negative selection using the MojoSort Human CD4+ T Cell Isolation kit (Biolegend) according to the manufacturer’s instructions. CD45RA+ve naïve and CD45RA-ve memory populations were further separated after CD4+ T cell isolation with CD45RA MicroBeads (Biolegend). For activated CD4+ T cells, PBMCs were treated with 5μg/ml PHA (Sigma) and 10IU/ml IL2 (Centre For AIDS Reagents, National Institute of Biological Standards and Control, UK [CFAR]) in RPMI1640 with 20% FBS for 72h prior to CD4+ T cell isolation. Once purified, CD4+ T cells were cultured in RPMI supplemented with 20% FBS and 10IU/ml IL2. Jurkat T cell lines (Clone E6-1; ATCC TIB-152) were cultured in RPMI with 10% FBS and 100U/ml penicillin/streptomycin. HEK 293T/17 cells (ATCC, CRL-11268) were cultured in DMEM with 10% FBS and 100U/ml penicillin/streptomycin. Tonsil tissue was obtained from an individual with primary HIV infection who underwent routine tonsillectomy (2 months after commencement of ART). As previously described^46^,the tonsillar tissue from elective tonsillectomy was dissected and mechanically digested, prior to cryopreservation of the cellular suspension. This was collected under the Imperial College Infectious Diseases Biobank (REC: 15/SC/0089). Lymph nodes were obtained from the field of surgery of participants undergoing surgery for diagnostic purposes and/or complications of inflammatory lung disease. Informed consent was obtained from each participant, and the study protocol approved by the University of KwaZulu-Natal Institutional Review Board (approval BE024/09).

### Plasmids and virus production

The HIV-1 clone pNL4.3 was obtained from the CFAR, NIBSC (cat# 2006). HIV-1 NL4.3 ΔNef and pNL4.3 ΔVpr were provided by R. Sloan (University of Edinburgh, UK)^47^. NL4.3 ΔVpu was provided by S. Neil (King’s College London, UK)^48^. NLENG1-IRES was provided by D. Levy (NYU, USA)^49^. NL4.3 bearing the CCR5-tropic BaL *Env* was provided by G. Towers (UCL, UK)^50^. CCR5 tropic transmitter/founder virus plasmids CH044 and CH077 were provided by G. Towers (UCL, UK) and were originally obtained through the NIH AIDS Reagent Program [NIHARP], Division of AIDS, NIAID, NIH: pCH040.c/2625 (cat# 11740) and pCH077.t/2627 (cat# 11742) from Dr. John Kappes and Dr. Christina Ochsenbauer. NL4.3 Vpr Q65R, NL4.3 Vpr S79A, NL4.3 Vpr R80A were generated by site-directed mutagenesis (Promega) using the following primers:

NL4.3 VprQ65R fw:GTGGAAGCCATAATAAGAATTCTGCGACAACTGCTGTTTATCCATTTCAG
NL4.3 VprQ65R rv:CTGAAATGGATAAACAGCAGTTGTCGCAGAATTCTTATTATGGCTTCCAC
NL4.3 Vpr S79A fw: GAATTGGGTGTCGACATGCCAGAATAGGCGTTACTC
NL4.3 Vpr S79A rv:GAGTAACGCCTATTCTGGCATGTCGACACCCAATTC
NL4.3 Vpr R80A fw: GGTGTCGACATAGCGCAATAGGCGTTACTCG
NL4.3 Vpr R80A rv: CGAGTAACGCCTATTGCGCTATGTCGACACC.

All virus stocks were produced by plasmid transfection of HEK 293T cells with Fugene 6 (Promega). Supernatants were harvested at 48h and 72h, filtered, DNase treated, purified and concentrated by ultracentrifugation through a 25% sucrose cushion and resuspended in RPMI1640 with 10% FBS. Viral titres were determined by measuring reverse transcriptase activity by SG-PERT assay^51^.

### HIV-1 infection and cell-to-cell spread

For cell-to-cell spread experiments, activated primary CD4+ T cells (donor cells) were infected with 800mU reverse transcriptase per 10^6^ cells of HIV-1 by spinoculation at 1200x*g* for 2h at room temperature and incubated in RPMI 20% FBS supplemented with 10IU/ml IL2 for 72h. HIV-1+ donor CD4+ T cells were washed with medium, counted and cultured with primary CD4+ target T cells at a 1:1 ratio in RPMI 20% FBS supplemented with 10IU/ml IL2 for up to 72h before analysis by flow cytometry or FACS sorting. Uninfected target CD4+ T cells were pre-stained with 1-2nM CellTrace FarRed dye (Invitrogen) prior to co-culture. For cell-to-cell spread into tonsil-derived lymphocytes, total tonsil lymphocytes were cultured at a 4:1 ratio with HIV-1 infected or uninfected eFluor450-labelled Jurkat T cells. For FACS sorting experiments, donor cells were pre-labeled with cell dye eFluor450 (ThermoFisher). For transwell experiments, HIV-1 infected donor T cells were separated from target T cells by a 0.4μm transwell insert (Corning). Experiments to quantify cell-to-cell versus cell-free infection in the presence and absence of a transwell were performed in equivalent volumes (600μl). For some experiments, FACS sorted infected resting CD4+ target T cells were returned into cultured for up to 4 days. Infection levels were measured by intracellular Gag staining and flow cytometry, and virus release into cell culture supernatant determined by SG-PERT^51^. At day 1 or day 4 post FACS sorting, resting CD4+ T cells were washed extensively and co-cultured at a 1:1 ratio with uninfected eFluor450-labelled Jurkat T cells for 72h, when Jurkat T cell infection was measured by Gag-staining. Where indicated, cultures were incubated in the presence of 20ng/ml IL7 (Miltenyi Biotec), 20ng/ml IL15 (Peprotech), 20ng/ml IL12 (Peprotech) or 50ng/ml TGFβ (Peprotech). The following inhibitors were added 30min before co-culture at the following concentrations: T20 (25-50ng/ml, CFAR), Efavirenz (1μM, CFAR), Raltegravir (5μM, CFAR) and Ruxolitinib (50nM, Sigma). For RNAi knockdown of DCAF1, primary CD4+T cells were activated for 4 days with 1μg/ml plate-bound αCD3 antibody (cloneOKT3, Biolegend) in the presence of 2 μg/mL soluble αCD28 antibody(clone CD28.2, Biolegend). RNAi knockdown of DCAF1 was performed as described before^52^ using ON-TARGET plusHuman DCAF1 siRNA - SMARTpool (Dharmacon) and non-targeting siRNA (Dharmacon) was used as a control. 2×10^6^cells were electroporated with 200 pmol siRNA using a NeonTransfection System (Thermo Fisher Scientific; three pulses, 10 ms, 1,600 V). After 48h, DCAF1 knockdowns were confirmed by western blotting and cells used in cell-to-cell spread experiments as described above.

### Flow cytometry and FACS

For flow cytometry analysis, cells were washed in PBS and stained with fixable Zombie UV Live/Dead dye, Aqua Live/Dead dye or NIR Live/Dead dye (Biolegend) for 5 min at 37°C. Excess stain was quenched with FBS-complemented RPMI. When tonsil and lymph node lymphocytes were used, Live/Dead staining was quenched using human AB serum (Sigma) in RPMI. Cell surface staining was performed in PBS, complemented with 20% Super Bright Staining Buffer (ThermoFisher) when appropriate, at 4°C for 30min. Unbound antibody was washed off thoroughly and cells were fixed with 4% FA or PFA before intracellular staining. For intracellular detection of IFNγ in infected target CD4+ T cells after 72h of cell-to-cell spread, cells were treated throughout the co-culture with IL7 and Brefeldin A (Biolegend) treated for 6h before surface staining and fixation. Permeabilisation for intracellular staining was performed with IC perm buffer or FoxP3 Buffer set (Biolegend) according to the manufacturer’s instructions. For detection of intracellular P-STAT5, cells were resuspended in ice cold True-Phos Perm buffer (Biolegend) and permeabilised for 48h at -20°C. Intracellular P-STAT5 staining was then performed in PBS with wash steps performed at 1800 rpm for 6 min at 4°C.The following antibody clones and fluorochromes were used: CD3 (UCHT1, Biolegend; Bv510, Bv711, FITC), CD8 (SK1, Biolegend; Bv605, PE), CD4 (SK3, Biolegend; APC/Fire750); CD45RA (HI100, Biolegend; Bv421, PE-Dazzle); CD45RO (UCHL1, Biolegend; PerCp-Cy5.5), CD69 (FN50, Biolegend; APC/Fire750, PE-Dazzle); CXCR6/CD186 (K041ES, Biolegend; PE-Dazzle); MCM2 (ab4461, ABCAM; was detected with a secondary anti-rabbit AlexaFluor488-tagged antibody); HLA-DR (L234, Biolegend; PerCp-Cy5.5); CD49a (TS2/7, Biolegend; PE-Cy7); PD-1 (EH12.2H7, Biolegend; PE-Cy7); Ki67 (Ki-67, Biolegend; Bv711, PE); Blimp-1 (6D3, BD Pharmingen; PE); CD101 (BB27, Biolegend; PE-Cy7); CX3CR1 (2A9-1, Biolegend; PE-Dazzle); CD103 (Ber-ACT8, Biolegend; Bv711); CD127 (AO19D5, Biolegend; PE-Cy7), IFNγ (B27, Biolegend; PE), Phospho-STAT5 (Clone 47/Stat5 (pY694), BD; PE) and HIV-1 Gag core antigen (FH190-1-1, Beckman Coulter; PE, FITC). All samples were acquired on either an BD Fortessa X20 or LSR II using BD FACSDiva software and analyzed using FlowJo v10 (Tree Star). FACS sorting was performed with a BD FACSAria III or BD FACSAria IIu Cell Sorter. Cells were either lysed immediately in RLT lysis buffer (Qiagen) with 1% β-mercaptoethanol (Sigma) and stored at -80C for later RNA extraction or resuspended in RPMI supplemented with 20% FBS and 10IU/ml IL2 and used immediately.

### Western blotting

Virus-containing supernatants (normalised for equal loading by measuring RT activity) or fifteen micrograms of total CD4+ T cell protein lysate were separated by SDS-PAGE, transferred onto nitrocellulose and blocked in PBS with 0.05% Tween 20 (v/v) and 5% skimmed milk (w/v). Blots were probed with rabbit antisera raised against HIV-1 Gag p24 (cat# 0432 donated by Dr G. Reid and obtained from the CFAR), Vpr anti-serum (cat# 3951, NIH ARP), α−alpha-Tubulin (T6199, Sigma-Aldrich) and α -DCAF1 antibody (11612-1-AO, Proteintech), followed by goat anti-rabbit or goat anti-mouse IRdye 800CW or 680RD infrared secondary antibody (Abcam) and imaged using an Odyssey Infrared Imager (LI-COR Biosciences) and analysed with Image Studio Lite software.

### Quantification of HIV-1 integration

To quantify integration of HIV-1 in resting T cells, nested Alu-gag quantitative PCR was performed as previously described^53^. Briefly, DNA was isolated from FACS sorted infected resting CD4+ memory T cells after 72h of cell-to-cell spread using the Qiagen Blood Mini Kit. Integrated DNA was pre-amplified using 100nM Alu fw primer, 600nM HIV-1 Gag rv primer, 0.2mM dNTP, 1U Phusion Hot Start Flex (Promega), and 45ng DNA in 50μl reactions. Cycling conditions were: 94°C for 30s, followed by 40 cycles of 94°C for 10s, 55°C for 30s, and 70°C for 2.5min. For quantitation of HIV-1 integration, a second round real-time quantitative PCR was performed using the pre-amplified DNA. These samples were run alongside a standard curve of known dilutions of CEM cells containing integrated HIV-1 DNA. Reactions contained 0.25μM of RU5 fw and rv primers, and 0.2μM probes, 1x Qiagen Multiplex Mastermix, and 10μl pre-amplified DNA. Cyclin conditions were: 95°C for 15min, followed by 50 cycles of 94°C for 60s and 60°C for 60s. 2LTR circles were measured by quantitative PCR^54^. Reactions contained 150ng DNA, 10μ 2LTR fw and rv primers, 10μM probe and 1x TaqMan Gene Expression Master Mix (ThermoFisher). Cycling conditions were: 95°C for 15min, followed by 50 cycles of 95°C for 15s and 60°C for 90s. Reactions were performed using 7500 Real-Time PCR System (Applied Biosystems). The following primers and probes were used:

Alu fw: GCCTCCCAAAGTGCTGGGATTACAG
HIV-1 Gag rv: GTTCCTGCTATGTCACTTCC
RU5 fw: TTAAGCCTCAATAAAGCTTGCC
RU5 rv: GTTCGGGCGCCACTGCTAGA
RU5-WT probe: FAM-CCAGAGTCACACAACAGACGGGCACA-TAMRA
RU5-degenerate 1 probe: FAM-CCAGAGTCACATAACAGACGGGCACA-TAMRA
RU5-degenerate 2 probe: FAM-CCAGAGTCACACAACAGATGGGCACA-TAMRA
2LTR fw: AACTAGAGATCCCTCAGACCCTTTT
2LTR rv: CTTGTCTTCGTTGGGAGTGAAT
2LTR probe: FAM-CTAGAGATTTTCCACACTGAC-TAMRA

### qRT-PCR

RNA was extracted from FACS sorted target memory CD4+ T cells with RNeasy Micro Kit (Qiagen) according to the manufacturer’s instructions. cDNA was synthesised using SuperScript IV with random hexamer primers (Invitrogen) and qRT-PCR was performed using Fast SYBR Green Master Mix and 7500 Real-Time PCR System (Applied Biosystems). Gene expression was determined using the 2^-ΔΔCt^ method and normalised to GAPDH expression. The following primers were used:

*GAPDH* fw: ACATCGCTCAGACACCATG, rv: TGTAGTTGAGGTCAATGAAGGG;
*CXCR6* fw: GACTATGGGTTCAGCAGTTTCA, rv:GGCTCTGCAACTTATGGTAGAAG;
*PRDM1* fw: ATGCGGATATGACTCTGTGGA, rv: CTGAACCGAAGTACCGCCATC;
*CD69* fw: ATTGTCCAGGCCAATACACATT, rv: CCTCTCTACCTGCGTATCGTTTT;
*S1PR1* fw: TCTGCTGGCAAATTCAAGCGA, rv: GTTGTCCCCTTCGTCTTTCTG;
*KLF2* fw: CTACACCAAGAGTTCGCATCTG; rv: CCGTGTGCTTTCGGTAGTG.

### Whole transcriptome profiling by RNA-Sequencing

RNA was extracted from FACS sorted target memory CD4+ T cells with RNeasy Micro Kit (Qiagen) according to the manufacturer’s instructions. For preparation of RNA-Sequencing libraries, RNA concentration was measured using the Qubit RNA High Sensitivity kit (Life Technologies) and quality checked on the 4200 Tapestation using either the High Sensitivity or standard RNA ScreenTape assay (Agilent Technologies), depending on the measured RNA concentrations. PolyA-tailed mRNA was separated for sequencing during library preparation. Libraries were prepared using KAPA’s mRNA HyperPrep kit (Roche Diagnostics) according to the manufacturer’s instructions using an input of up to 200ng and a fragmentation incubation time of 8 minutes at 94°C. Samples were sequenced on Illumina’s NextSeq500 (Illumina Cambridge) using a high output 75 cycle paired-end run. 24 libraries were multiplexed in the same run. Libraries were pooled in equimolar quantities, calculated from concentrations measured using the Qubit dsDNA High Sensitivity kit (Life Technologies) and fragment analysis using the D1000 High Sensitivity assay on the 4200 Tapestation (Agilent Technologies).

RNA sequencing data was quality assessed using FASTQC (https://www.bioinformatics.babraham.ac.uk/projects/fastqc/) before and after low-quality and adapter trimming using Trimmomatic^55^. Filtered reads were then pseudo-mapped using Kallisto^56^ to the transcriptome available in Ensembl v.101 (http://aug2020.archive.ensembl.org/index.html). Per-transcript counts were imported and aggregated per gene using the TXimport R package^57^. The DESeq2 package^58^ was used for data normalisation, outlier detection and differential gene expression analysis between biological groups. The DESeq2 results were ranked based on the log2 transformation of the adjusted p-values, to provide a pre-ranked list for Gene Set Enrichment Analysis (GSEA)^59^ as described in the GSEA documentation. Pathway enrichment and uptream regulator analysis was performed using Gene Set Enrichment Analysis (GSEA)^59^ and Ingenuity Pathway Analysis (IPA,) respectively. Heatmaps were generated using ClustVis (https://biit.cs.ut.ee/clustvis/) ^60^

### Transcriptomic comparison with published human T_RM_ cells

TPM data from previously published transcriptomes of human T_RM_ cells (GSE94964)^21^ were summed on gene level with Ensembl gene ID, gene name, and gene biotype using tximport and BioMart^57, 61^. TPM values <0.001 were adjusted to 0.001 as a lower limit of detection. These data were aligned to the transcriptomic data from the present study using gene symbol in an integrated Log_2_ transformed data matrix and subjected to batch correction by study using Combat^62^. Expression of selected genes previously identified to be up and downregulated in T_RM_ were used to cluster the samples in both studies using 1-Spearman rank correlation with average linkage in ClustVis^60^. A transcriptional signature score for T_RM_ was derived from the difference between the sum of up and down-regulated genes in T_RM_ in the previously published signature. This score was used to evaluate the relative similarity of each transcriptome data set in the present project to T_RM_ and non-T_RM_ data.

### Statistical analysis

Statistical analysis was performed using GraphPad Prism. Normally distributed data was analysed for statistical significance by two-tailed *t-*tests (when comparing two groups) or one-way ANOVA with Bonferroni or Dunnett’s post-test (when comparing more than two groups). Data show the mean +/- the S.E.M with significance shown on the figures. Where appropriate, the median+IQR is shown and Kruskall-Wallis test was used to compare groups. Significance levels were defined as *, P < 0.05; **, P < 0.01 and ***, P < 0.001.

## Acknowledgements

This work was funded by a Wellcome Trust Investigator award (108079/Z/15/Z) to C.J. We are grateful to members of the Jolly lab, as well as Greg Towers, Rebecca Sumner, Lorena Zulianai - Alvarez, Lucy Thorne, Laura McCoy and Richard Milne for helpful discussions and critical reading of the manuscript. We thank Jamie Evans (UCL) for assistance with sorting CD69-negative T cells and Parisa Amjadi (Imperial College London Chelsea and Westminster Hospital) for sorting target T cells from co-culture experiments and the Pathogen Genomics Unit (PGU) at UCL for RNAseq analysis. We acknowledge the NIBSC Centre for AIDS Reagents, the NIH AIDS Reagent Program and Imperial College NIHR Biomedical Research Centre, London, UK.

## Author contributions

A.K.R., and C.J. conceived the project. A.K.R., and C.J. designed the experiments. A.K.R., M.S. and D.M performed the experiments. A.K.R., C.J., L.P., M.K.M., M.S., D.M., A.G-A and M.N. analysed the data. L.P. and M.K.M provided reagents. J.P.T., C.H., S.F., R.M, K.J.D., and A.S. provided lymphoid tissue samples. A.G-A and M.N. performed the core T_RM_ transcriptional mapping analysis. A.K.R., and C.J. prepared the manuscript. All authors provided critical review of the manuscript.

---

**Extended data Fig. 1.**
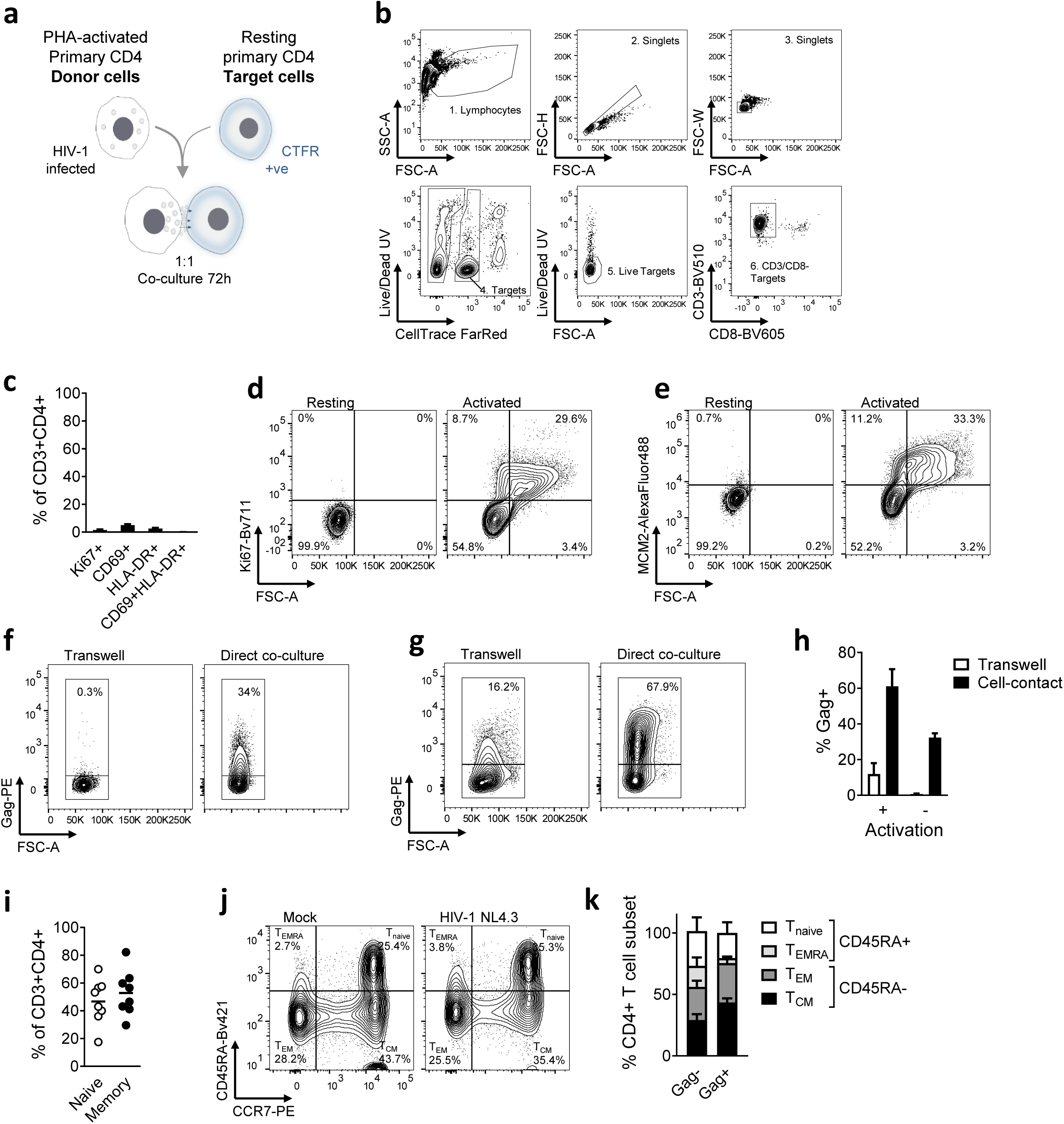
(**a**) Experimental set-up schematic. (**b**) Flow cytometry gating strategy. (**c**) Ki67, CD69 and HLA-DR expression of CD3+CD4+ T cells from unstimulated PBMCs (n=8). (**d**) Ki67 expression on resting and activated primary CD4+ T cells. Representative flow cytometry plots. (**e**) MCM2 expression on resting and activated primary CD4+ T cells. Representative flow cytometry plots. (**f**) Resting or (**g**) mitogenically-activated primary target CD4+ T cells cultured with HIV-1 infected Jurkat T cells separated by a 0.4μm transwell or in direct co-culture. Target cell infection levels was measured by intracellular staining for Gag. Representative flow cytometry plots are shown. (**h**) Infection levels of target CD4+ T cells determined by intracellular Gag staining and flow cytometry (n=2). (**i**) Proportion of CD45RA+ naive and CD45RA-memory CD4+ T cells in unstimulated PBMCs (n=8). (**j**) Resting target CD4+ T cells were cultured with mock-treated or HIV-1-infected donor cells. Surface expression of CD45RA and CCR7 were measured after 72h of co-culture. Representative flow cytometry plots are shown. (**k**) Quantification of T cell subsets in infected (Gag+) and uninfected (Gag-) resting CD4+ T cells according to CD45RA/CD62L expression (n=3). T_naive_ = CD45RA+/CD62L+, T_EMRA_ = CD45RA+/CD62L-, T_EM_ = CD45RA-/CD62L-, T_CM_ = CD45RA-/CD62L+. Data are shown as mean±SEM.

**Extended data Fig. 2.**
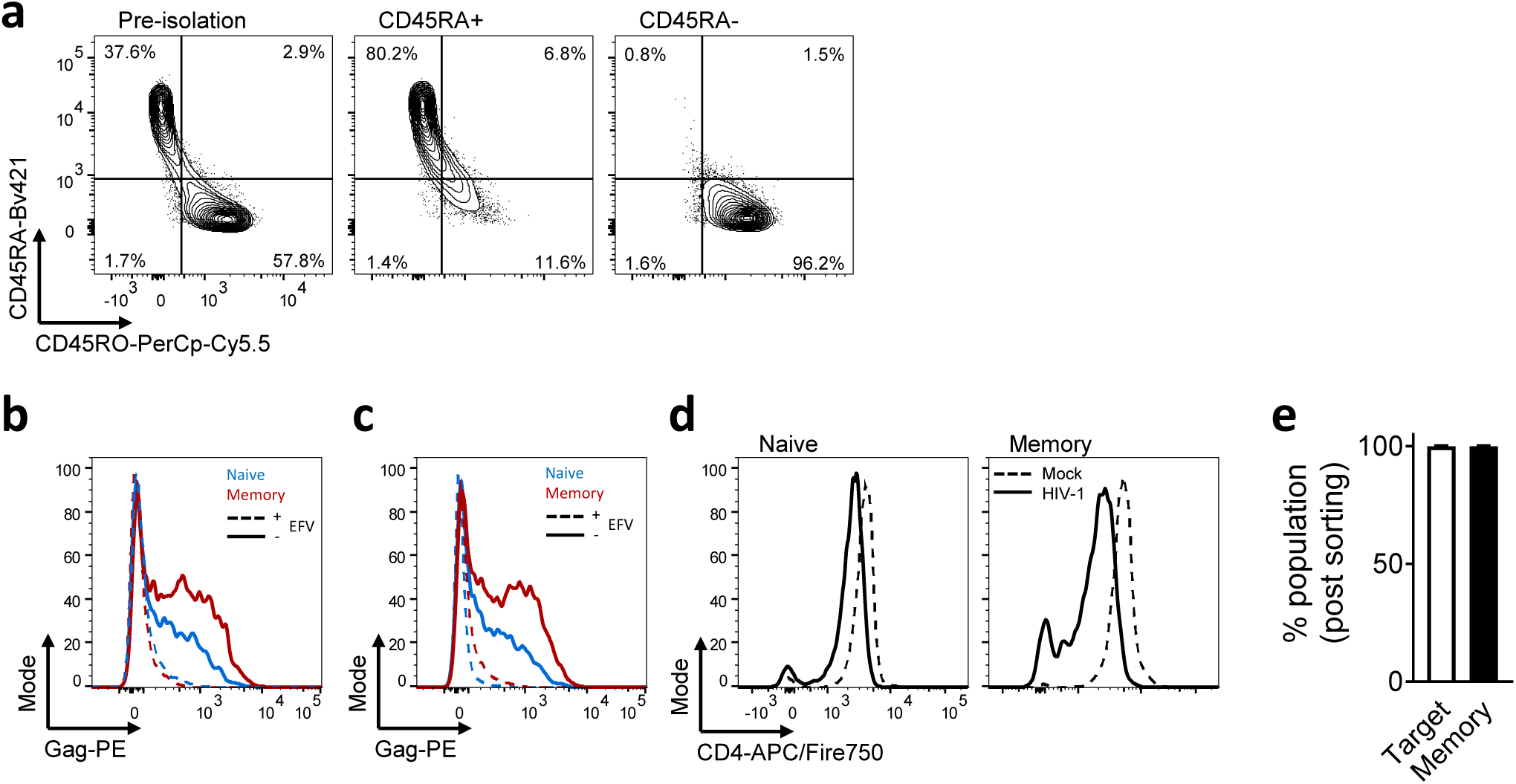
(**a**) Representative flow cytometry plots of CD45RA+ and CD45RA-CD4+ T cells pre- and post-isolation. (**b**) Representative histogram of intracellular Gag staining in resting naïve (CD45RA+) and memory (CD45RA-) CD4+ T cells after 72h of cell-to-cell spread ± T20. (**c**) Representative histogram of intracellular Gag-levels in resting naïve (CD45RA+) and memory (CD45RA-) CD4+ T cells after 72h of cell-to-cell spread ± Efavirenz. (**d**) Representative histogram of CD4 surface levels in resting naïve (CD45RA+) and memory (CD45RA-) CD3+ T cells after 72h of cell-to-cell spread. (**e**) Mean post-sort population purity of T cells from (Fig. 1 n-p) was 99.92% target cells of which 99.86% were memory T cells (n=5).

**Extended data Fig. 3.**
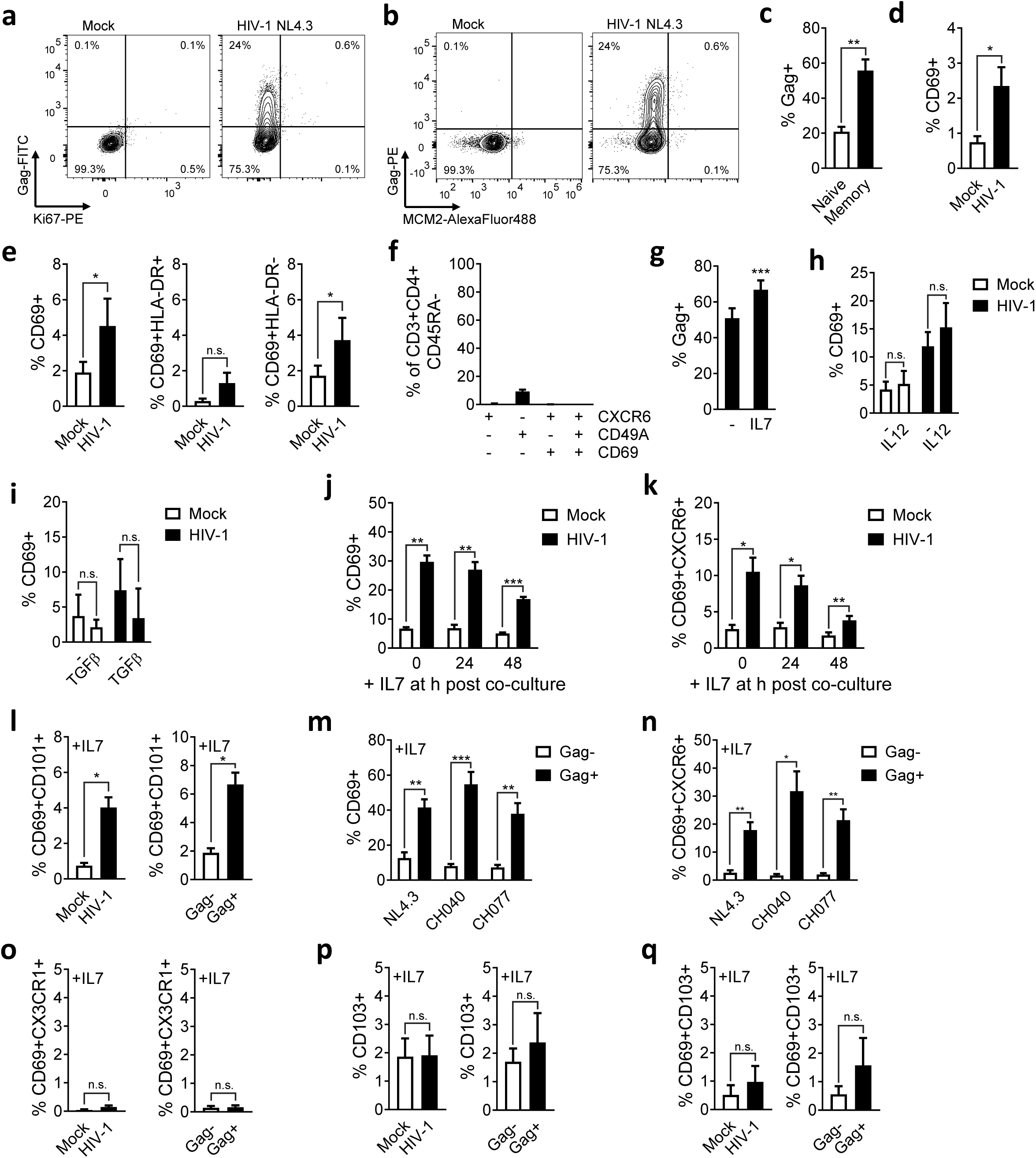
(**a** and **b**) Representative flow cytometry plots showing (**a**) Ki67 and Gag, or (**b**) MCM2 and Gag staining of resting CD4+ T cells after co-culture with mock or HIV-1 infected primary donor CD4+ T cells. (**c**) FACS sorted CD69- resting naïve or memory CD4+ T cells co-cultured with HIV-1 infected primary CD4+ donor T cells and infection of targets measured by Gag staining (n=4). (**d**) FACS sorted CD69- CD4+ T cells co-cultured with HIV-1 infected primary CD4+ donor T cells. CD69 expression was measured on resting memory CD4+ T cells (n=4). (**e**) Total CD69 expression alongside CD69 with or without HLA-DR co-expression on infected resting memory T cells (n=6). (**f**) Expression of T_RM_-markers CXCR6, CD49A and CD69 on resting CD4+ memory T cells from unstimulated PBMCs (n=8). (**g**) Quantification of cell-to-cell spread of HIV-1 WT to resting memory CD4+ T cells in the presence or absence of IL7 (n=10). (**h** and **i**) CD69 expression on infected resting memory CD4+ target T cells in the presence or absence of (**h**) IL12 (n=4) or (**i**) TFGβ (n=4). (**j**) CD69 expression on infected resting memory CD4+ T cells with IL7 added at the indicated times post cell-mixing (n=5). (**k**) CD69/CXCR6 co-expression from (**j**) (n=5). (**l**) CD69/CD101 co-expression on infected resting memory CD4+ T cells (n=3) (**m**) CD69 upregulation in response to IL7 on resting memory CD4+ T cells infected HIV-1 NL4.3, or transmitter-founder viruses CH040 and CH077 comparing infected Gag+ and uninfected Gag-bystander cells (n=7). (**n**) CD69/CXCR6 co-expression on resting memory CD4+ T cells from (**i**) (n=7). (**o**) CD69/CX3CR1 co-expression on infected resting memory CD4+ T cells (n=3). (**p**) CD103 expression and (**q**) CD69/CD103 co-expression on infected resting memory CD4+ T cells (n=4). Data are the mean±SEM. Paired two-tailed *t*-test or one-way ANOVA with Bonferroni post-test was used. For (**i**), median+IQR is shown and Kruskall-Wallis test was used to compare groups *, p<0.05; **, p<0.01; ***, p<0.001; n.s., not significant.

**Extended data Fig. 4.**
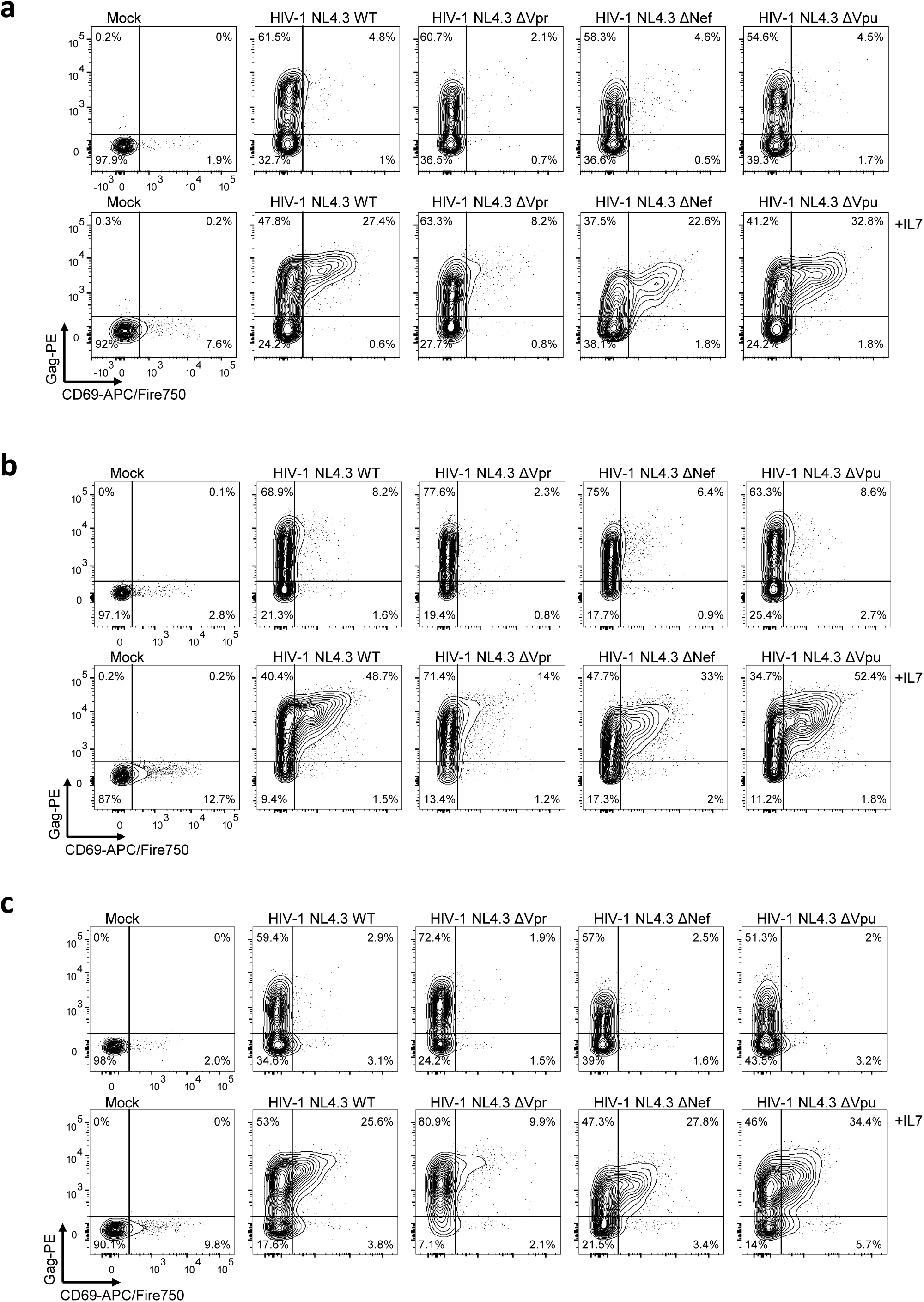
Resting memory CD4+ T cells were co-cultured with HIV-1 infected primary CD4+ T cells infected with HIV-1 WT or mutant viruses. Representative flow cytometry plots of HIV-1 Gag and CD69 co-expression in the presence or absence of IL7 from three independent experiments are shown (**a-c**).

**Extended data Fig. 5.**
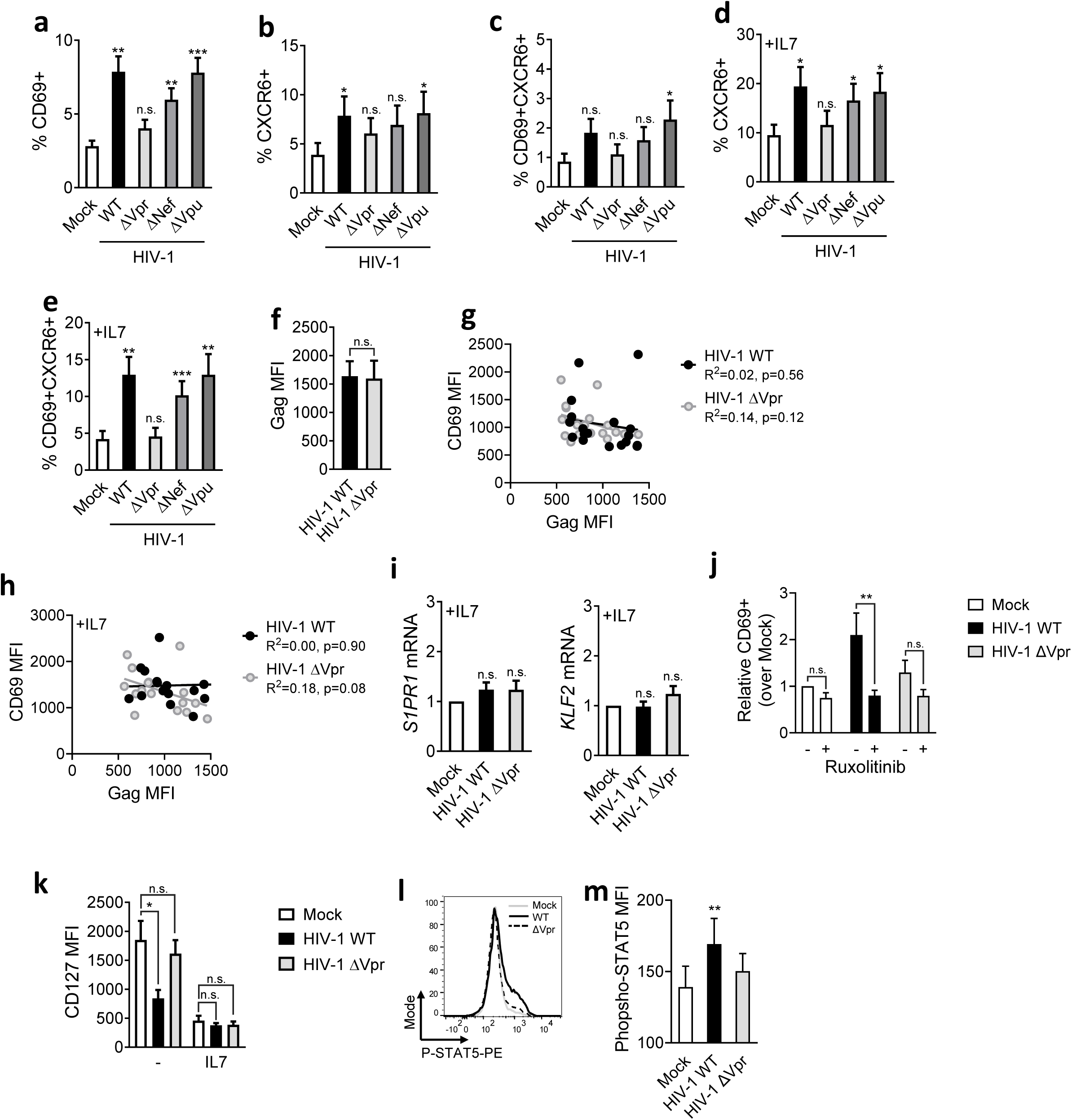
(**a**) CD69 surface expression on resting CD45RA- memory CD4+ T cells following co-culture with primary CD4+ donor T cells infected with HIV-1 NL4.3 (WT), ΔVpr, ΔNef or ΔVpu or uninfected (mock) donors (n=9). (**b**) CXCR6 expression from (**a**) (n=9). (**c**) CD69/CXCR6 co-expression from (**a**) (n=9). (**d**) As for (b) but cells were incubated in the presence of IL7 (n=9). (**e**) CD69/CXCR6 surface co-expression from (d) (n=9). (**f**) Gag MFI of cell-to-cell spread of HIV-1 WT and ΔVpr to resting memory CD4+ T cells (n=10). Correlation plot of CD69 MFI with Gag MFI in absence (**g**) or presence (**h**) of IL7 (n=18). (**i**) *S1PR1* and *KLF2* mRNA levels in FACS sorted resting memory CD4+ T cells from Fig. 3e. Fold change over mock is shown (n=5). (**j**) CD69 expression on infected resting memory CD4+ T cells ± Ruxilitinib (n=4). (**k**) CD127 MFI on infected resting memory CD4+ T cells ± IL7 (n=7). (**l**) Representative histogram of intracellular STAT5-phosphorylation in infected resting memory T cells. (**m**) Quantification of (**l**) (n=10). Data are the mean±SEM. One-way ANOVA with Dunnet’s post-test was used. Statistical significance is shown relative to mock treated cells (no HIV-1). R^2^ in (g) and (h) was determined by simple linear regression. *, p<0.05; **, p<0.01; ***, p<0.001; n.s., not significant.

**Extended data Fig. 6.**
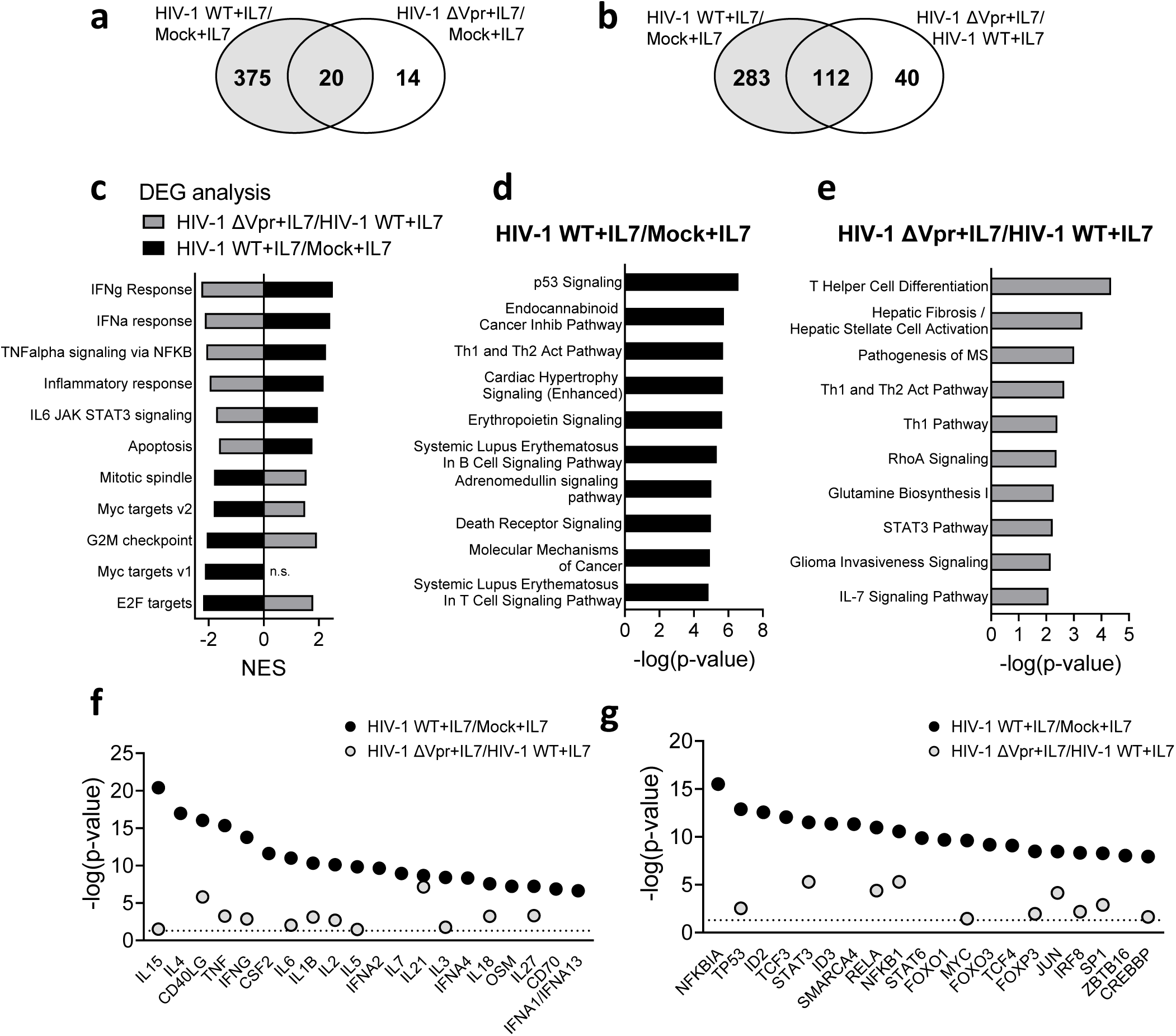
(**a**) and (**b**) Venn diagrams showing overlap of DEGs comparing expression profiles of HIV-1 WT+IL7/Mock+IL7 with (**e**) HIV-1 ΔVpr+IL7/\Mock+IL7 or (**f**) HIV-1 ΔVpr+IL7/HIV-1 WT+IL7. (**g**) GSEA was performed on expression profiles comparing HIV-1 WT+IL7/Mock+IL7 (black) or HIV-1 ΔVpr+IL7/HIV-1 WT+IL7 (grey). Normalised enrichment scores are shown for significantly enriched Hallmark gene sets are shown (FDR q-value<0.05 and NES>1.75). (**h**) and (**i**) top ten significantly enriched canonical pathways predicted by ingenuity pathway (IPA) analysis of DEGs (h) HIV-1 WT+IL7/Mock+IL7 or (i) HIV-1 ΔVpr+IL7/HIV-1 WT+IL7 (adjusted p-value<0.05). (**j**) Cytokines and (**k**) transcription regulators predicted to be upstream regulators by IPA of gene expression signatures HIV-1 WT+IL7/Mock+IL7 (black) or HIV-1 ΔVpr+IL7/Mock+IL7 (grey), line indicates p=0.05.

**Extended data Fig. 7.**
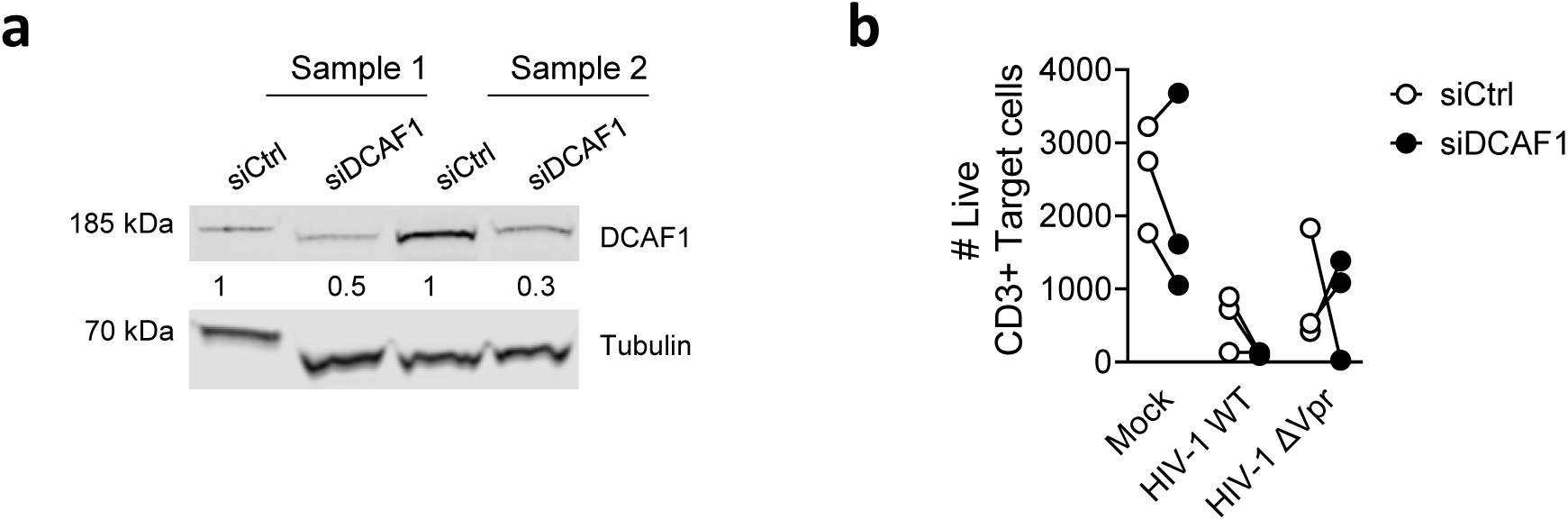
(**a**) Western blot showing siRNA knockdown of DCAF1 in CD3/CD28-activated CD4+ T cells 48h post transfection. Two representative samples are shown. (**b**) Number of live CD3+ Target T cells recovered after 72h of cell-to-cell spread into control or DCAF1 siRNA-treated T cells (n=3).

